# Differential sorting behavior for soluble and transmembrane cargoes at the trans-Golgi network in endocrine cells

**DOI:** 10.1101/797134

**Authors:** Blake H. Hummer, Drew Maslar, Margarita Soltero Gutierrez, Noah F. de Leeuw, Cedric S. Asensio

## Abstract

Regulated secretion of neuropeptides and peptide hormones by secretory granules (SGs) is central to physiology. Formation of SGs occurs at the trans-Golgi network (TGN) where their soluble cargo aggregates to form a dense core, but the mechanisms controlling the sorting of regulated secretory cargoes (soluble and transmembrane) away from constitutively secreted proteins remain unclear. Optimizing the use of the retention using selective hooks (RUSH) method in (neuro-)endocrine cells, we now quantify TGN budding kinetics of constitutive and regulated secretory cargoes. We further show that, by monitoring two cargoes simultaneously, it becomes possible to visualize sorting to the constitutive and regulated secretory pathways in real-time. Further analysis of the localization of SG cargoes immediately after budding from the TGN revealed that, surprisingly, the bulk of two studied transmembrane SG cargoes (phogrin and VMAT2) does not sort directly onto SGs during budding, but rather exit the TGN into non-regulated vesicles to get incorporated to SGs at a later step. This differential behavior of soluble and transmembrane cargoes suggests a more complex model of SG biogenesis than anticipated.

## Introduction

Secretion of neuropeptides and peptide hormones such as insulin depends on their efficient sorting into secretory granules (SGs) that are capable of regulated exocytosis in response to an extracellular, physiological stimulus. Disturbances in their secretion underlie the development of pathological states such as type II diabetes. SGs bud from the trans-Golgi network (TGN) where their soluble cargoes aggregate to form a dense core (Borgonovo et al., 2006, Hou et al., 2009, Park et al., 2009, Tooze, 1998). Two mechanisms have been proposed for sorting to SGs (Arvan and Halban, 2004). In the sorting by retention model, interactions in the lumen of the TGN drive the formation of SGs. Consistent with this, SGs contain large amount of granulogenic proteins such as chromogranins that aggregate under the specific pH and redox conditions of the TGN. In this model, sorting to SGs thus occurs by default, with proteins destined for other organelles being removed from nascent SGs after budding from the TGN. This maturation can last up to several hours after budding and depends on clathrin as well as the adaptor protein AP-1 (Dittie et al., 1996, Dittie et al., 1997), but the functional significance of this process remains unclear and is likely to differ between specialized secretory cells. In contrast, the sorting for entry model involves active sorting of proteins destined for SGs at the TGN. In support of this model, biochemical analysis of budding from the TGN has demonstrated specific sorting of soluble proteins into SGs (Tooze and Huttner, 1990).

In addition to soluble cargoes, the membrane of SGs also contains several transmembrane proteins, and many of them such as the enzyme peptidylglycine α-amidating monooxygenase (PAM) and the endothelial adhesion molecule P-selectin depend on cytosolic sequences for their sorting to SGs (Blagoveshchenskaya et al., 1999, El Meskini et al., 2001, Milgram et al., 1996). The neuronal vesicular monoamine transporter (VMAT2) fills secretory vesicles with monoamine neurotransmitters for exocytotic release, and localizes preferentially to SGs in rat neuroendocrine PC12 cells (Liu et al., 1994, Weihe et al., 1994). The sorting of VMAT2 to SGs depends on a C-terminal cytoplasmic di-leucine-like motif with upstream acidic residues. Mutation of this motif diverts VMAT2 from the regulated to the constitutive secretory pathway, suggesting that these residues act as a sorting motif for the RSP (Krantz et al., 2000, Li et al., 2005). In addition, the other SG membrane proteins IA-2 and phogrin contain a similar sorting sequence, and phogrin also depends on this motif for sorting to the RSP in insulin-secreting cells (Torii et al., 2005). However, the complete mechanisms contributing to sorting to SGs and indeed SG production remain poorly understood.

One of the main limitations to address this problem comes from the lack of sensitive approaches to evaluate sorting into pathways originating at the TGN. In the past, an elegant budding assay that relies on protein sulfation at the TGN has been used to demonstrate the specific sorting of proteins into nascent SGs (Tooze and Huttner, 1990). However, this assay is limited to the analysis of soluble SG cargo proteins that are both highly abundant and sulfated. It has therefore not been possible to monitor emergence from the TGN of lower abundance proteins, in particular integral membrane proteins, that are presumably not sulfated.

Here we report the optimization of the retention using selective hook (RUSH) system (Boncompain et al., 2012) to visualize SG formation from (neuro-)endocrine cells and to obtain quantitative measurements of TGN budding of SG and non-SG soluble and transmembrane cargoes. We further monitor two cargoes simultaneously to visualize sorting to the constitutive and regulated secretory pathways in real-time. We demonstrate that the transmembrane SG markers phogrin and VMAT2 do not localize with insulin directly after leaving the TGN but rather get added to SGs after budding.

## Results and discussion

### Visualization of SG formation using RUSH

In order to synchronize and visualize the movement of cargoes along the secretory pathway in (neuro-)endocrine cells, we used the RUSH system (Boncompain et al., 2012). This approach relies on the compartmentalized expression of streptavidin acting as a “hook” for a co-expressed cargo fused to streptavidin-binding peptide (SBP). Exposure of cells to biotin leads to the rapid, synchronized release of the hooked cargoes. By using fluorescently tagged reporters, it becomes possible to follow a wave of cargo as it traffics through the secretory pathway. To test whether this approach could be applied to study the formation of SGs from (neuro-)endocrine cells, we began by determining the behavior of soluble SG cargoes. For this, we relied on neuropeptide Y (NPY) and secretogranin II (SgII) as their respective fusion to emdGFP (or other fluorescent proteins) targets properly to SGs in various (neuro-)endocrine cells including INS-1 and PC12 cells that we are using for this study (Gandasi et al., 2015, Taraska et al., 2003, Courel et al., 2008). We fused emdGFP-SBP to the C-terminus of NPY (**Fig. S1A**) and transfected INS-1 and PC12 cells with optimized hook and cargo (NPY-emdGFP-SBP) plasmids together with a TGN marker (sialyltransferase-TagRFP657). As a first step, we validated that our cargo trafficked properly through the secretory pathway. In absence of biotin, the reporter was found diffused across the cell, consistent with an ER localization, and treatment of cells with biotin led to its redistribution as expected. At 30 min, we observed strong localization at the TGN and following that time point the majority of the signal could be found in post-TGN vesicular carriers (**Fig. 1A, Fig. S1B**). After 1h of biotin treatment, NPY-emdGFP-SBP accumulated in discrete punctate structures that co-localized with endogenous insulin in INS-1 cells or endogenous SgII in PC12 cells (**Fig. 1B, Fig. S1C)**. Similarly, SgII-emdGFP-SBP (**Fig. S1A**) co-localized with insulin in INS-1 cells (**Fig. S1D**). We also determined the steady-state distribution of the reporters after incubating our cells with biotin for 24h. NPY-emdGFP-SBP accumulated in SGs found at the tip of processes and colocalized with insulin (for INS-1 cells) or SgII (for PC12 cells) and SgII-emdGFP-SBP colocalized with insulin in INS-1 cells (**Fig. 1B, Fig. S1C, D**). We also confirmed that NPY-emdGFP-SBP expressed as a full-length protein in both INS-1 and PC12 cells by western blot and found lower molecular bands in INS-1 cells indicative of processing (**Fig. S1E**). These data suggest that these professional secretory cells are able to sort cargoes efficiently even under artificial conditions leading to a large wave of cargoes.

**Figure 1.**
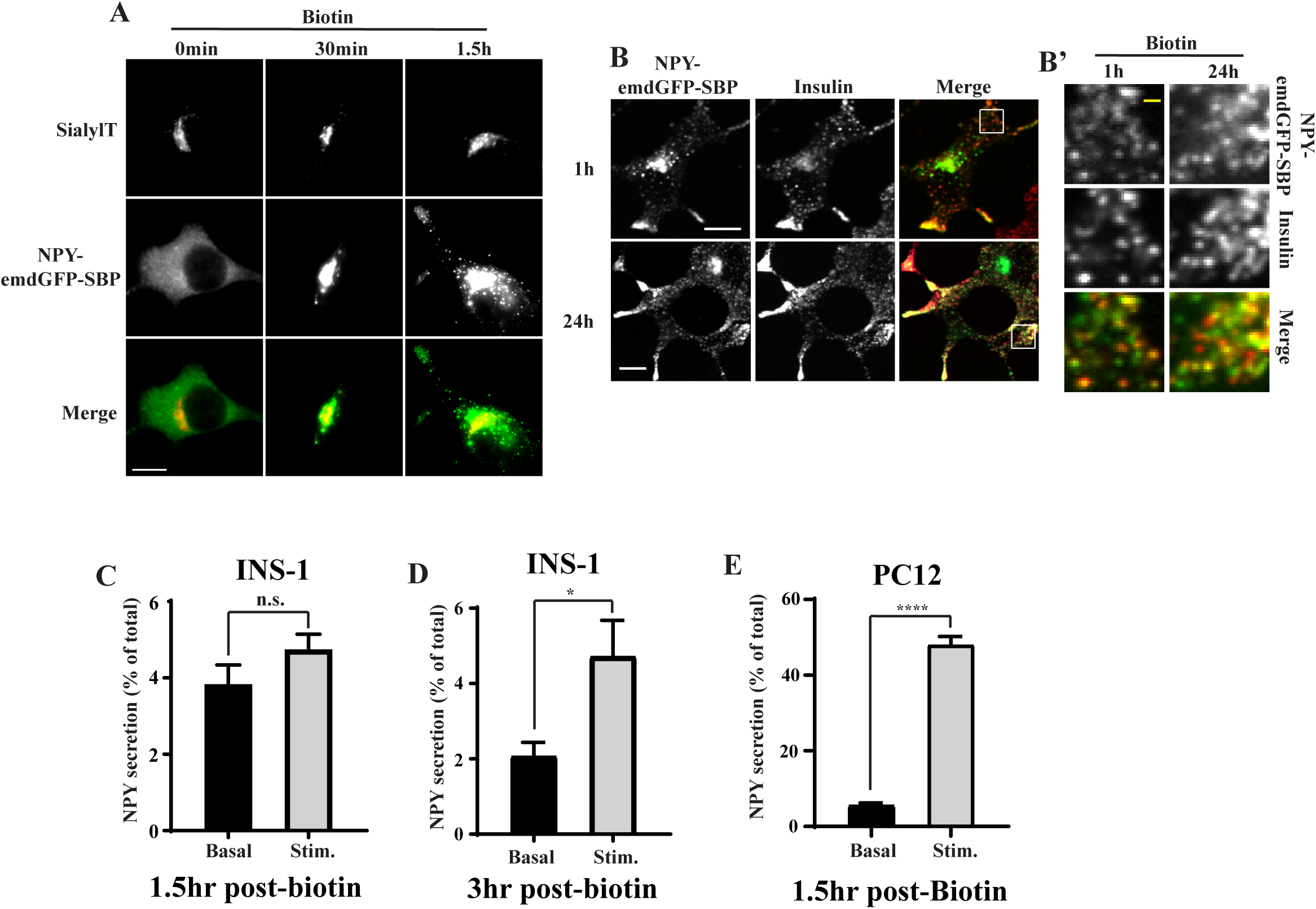
(**A**) INS-1 cells were transfected with NPY-emdGFP-SBP together with the TGN marker sialyltransferase-TagRFP657 and ER-hook, incubated with biotin for the indicated times, fixed and imaged using a widefield epifluorescence microscope. (**B**) INS-1 cells transfected with RUSH constructs were incubated with biotin for 1h or 24h, fixed and immunostained for insulin followed by Alexa Fluor 647-conjugated secondary antibodies. Cells were imaged by spinning-disk confocal microscopy. Insets are shown in **B’.**INS-1 (**C, D**) or PC12 cells (**E**) were transfected with NPY-emdGFP-SBP and ER hook for secretion assays. Cells were incubated with biotin for 1.5h (**C, E**) or 3h (**D**) followed by exposure to basal or stimulated conditions for 15min. Secretion was determined by reading the fluorescence in the media and was normalized to total fluorescence in cell lysate. *p < 0.05, ****p < 0.0001 relative to NPY basal (for INS-1 cells 1.5h n = 3, for INS-1 cells 3h n = 4, for PC12 cells n =3) by unpaired two-tailed t-test. Data shown indicate mean ± SEM. Scale bars indicate 10μm and 1μm for insets.

The ability to store SG cargo and release it under stimulatory conditions is a defining feature of (neuro-)endocrine cells. To further assess the functionality of our cargoes, we performed a secretion assay following RUSH. Working with INS-1 cells at first, we tested whether we could stimulate the release of NPY-emdGFP-SBP from newly formed SGs. Although we did not observe significant stimulation at 1.5h post biotin (**Fig. 1C**), the amount of NPY-emdGFP-SBP and SgII-emdGFP-SBP secreted in response to depolarization (high K+) more than doubled at 3h post biotin (**Fig. 1D**). We next performed experiments in PC12 cells and found that, under basal conditions, these cells secreted ~5% of total NPY-emdGFP-SBP; depolarization stimulated the release of NPY-emdGFP-SBP (~9 fold) at 1.5h post biotin (**Fig. 1E**). Altogether these data suggest that our cargoes sort efficiently to the regulated secretory pathway. The data also indicate that SGs from PC12 cells become competent for regulated release faster than insulin SGs from INS-1 cells.

### Determination of soluble and transmembrane SG cargo TGN budding kinetics

In order to visualize cargo movement along the secretory pathway, we imaged INS-1 cells expressing NPY-emdGFP-SBP or SgII-emdGFP-SBP live following the addition of biotin and observed a wave of fluorescent cargo entering the TGN labeled with sialyltransferase-TagRFP657. After reaching a peak, TGN fluorescence decreased concomitantly with the formation of post-TGN carriers (**Movie S1**). We determined the ER exit rate constant (k_ER_) by measuring the fluorescence change within the ER and fitting the data to a first-order decay curve. Next, we monitored the decrease in fluorescence at the TGN and fitted the data to extrapolate the TGN exit rate constant (k_TGN_). Both constructs exited the ER at similar rates but NPY-emdGFP-SBP exited the TGN slightly faster than SgII-emdGFP-SBP (**Fig. 2**). These data suggest that the trafficking rate through different organelles varies between cargoes. Consistent with this idea, the regulated secretory cargoes amylase and chymotrypsinogen undergo their main concentration between the ER and cis-Golgi in pancreatic exocrine cells as opposed to the TGN (Oprins et al., 2001). In the future, it will be interesting to compare the behavior of a variety of regulated secretory cargoes in different (neuro-)endocrine cells.

**Figure 2.**
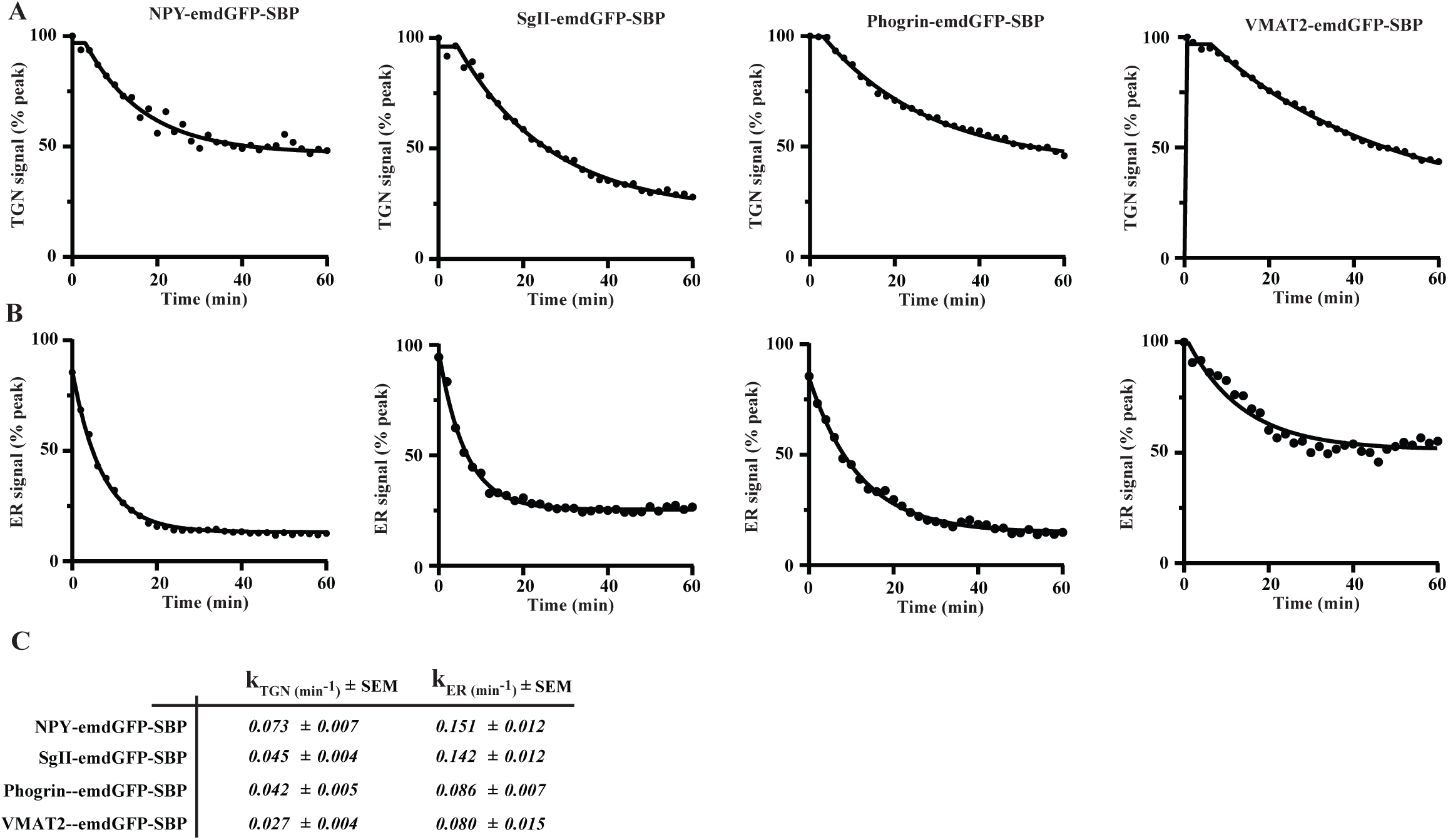
INS-1 cells were transfected as in **Fig. 1** with the indicated cargoes. Fluorescence within the TGN region (**A**) and the ER (**B**) was monitored and the data were fitted using a first-order kinetic model to extrapolate ER (k_ER_) and TGN rate constants (**C**). Representative curves are shown for each cargo. (n=16 for NPY-emdGFP-SBP from 3 independent transfections; n=8 for SgII-emdGFP-SBP from 2 independent transfections; n=14 for phogrin-emdGFP-SBP from 3 independent transfections; n=11 for VMAT2-emdGFP-SBP from 2 independent transfections).

Next, we tested whether RUSH could be applied to study budding kinetics of SG transmembrane protein cargoes. For this, we relied on phogrin and VMAT2 (**Fig. S1A**), which both target efficiently to SGs in both INS-1 and PC12 cells (Taraska et al., 2003, Tsuboi et al., 2000, Asensio et al., 2010, Liu et al., 1994). By western blot, we found that phogrin-emdGFP-SBP expressed as a full-length protein in both INS-1 and PC12 cells and observed lower molecular bands in INS-1 cells indicative of processing (**Fig. S1D**). In absence of biotin, phogrin-emdGFP-SBP was found diffused across the cell, consistent with an ER localization, and treatment of cells with biotin led to its redistribution as expected. At 40 min, we observed strong localization at the TGN and following that time point the majority of the signal could be found in post-TGN vesicular carriers (**Fig. S2A**). Live imaging experiments in INS-1 cells revealed that phogrin-emdGFP-SBP and VMAT2-emdGFP-SBP exited the ER with similar kinetics, which were slower than the ones measured for soluble regulated SG cargoes. Phogrin-emdGFP-SBP exited the TGN with similar kinetics than SgII-emdGFP-SBP, and VMAT2-emdGFP-SBP budding rate was significantly slower (**Fig. 2, Movie S1**). Importantly, our imaging conditions led to minimal photobleaching (**Fig. S2B, C**). Our data illustrate that secretory cargoes exit the TGN at specific rates with soluble SG cargoes tending to traffic faster than transmembrane SG cargoes. Intuitively, this seems logical as the relative enrichment of soluble vs transmembrane proteins on SGs is biased towards soluble cargoes by at least one or two orders of magnitude (Suckale and Solimena, 2010). It is also possible that these cargoes get modified to various extents as they traffic through the secretory pathway and that these protein modification steps contribute to the difference in kinetics.

### Transmembrane SG cargoes do not exit the TGN within SGs but get recruited later

Surprisingly, when looking at phogrin-emdGFP-SBP and VMAT2-emdGFP-SBP localization after 1h of biotin, we found that they did not colocalize with insulin unlike NPY-emdGFP-SBP and SgII-emdGFP-SBP (**Fig. 3**). However, we observed robust accumulation at the tip of processes after 24 h of biotin treatment, suggesting that these fusion proteins eventually traffic properly (**Fig. 3**). Time course experiments revealed that the enrichment within insulin granules increases gradually to peak at around 8h post biotin (**Fig. S2D)**. In addition, we obtained similar results after pulsing our cells with biotin for 30min followed by extensive washes to enable hooking of cargo synthesized after the initial wave (**Fig. S2E**). This indicates that continuous synthesis of cargo happening after the initial release of the hooked cargo is not responsible for the increase in colocalization observed at later time points. These data suggest that, at least using this approach, phogrin-emdGFP-SBP and VMAT2-emdGFP-SBP might exit the TGN within non-regulated vesicles, likely constitutive secretory carriers, and get incorporated to SGs only later, possibly during SG maturation.

**Figure 3.**
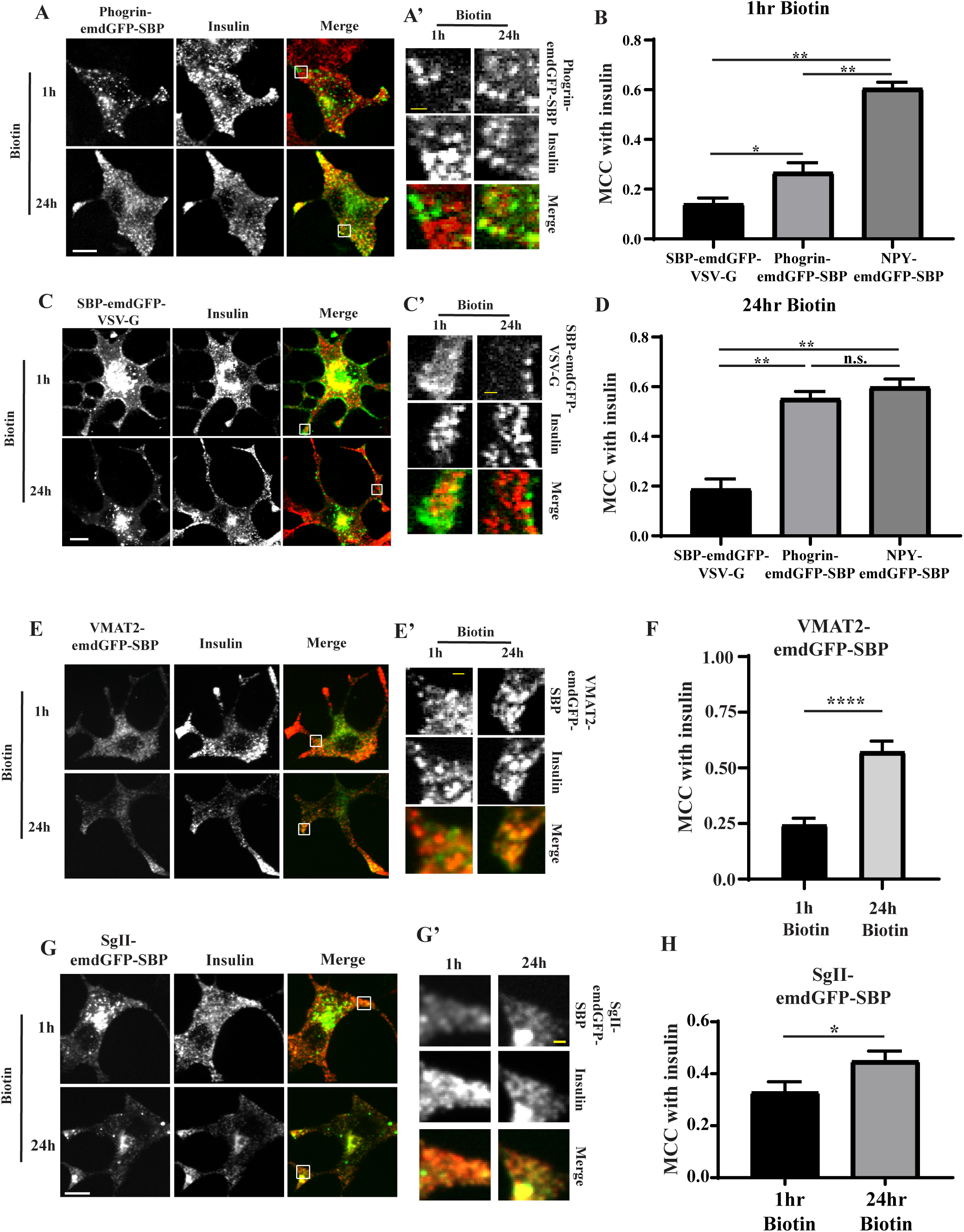
INS-1 cells were transfected with phogrin-emdGFP-SBP (**A**) or SBP-emdGFP-VSV-G (**B**) and incubated with biotin as in **Fig. 1**. (**A**) INS-1 cells were co-stained for insulin at 1h or 24h. Insets are shown in **A’** and **C’**. The extent of colocalization with insulin was determined by Manders Correlation Coefficient (MCC) at 1h (**B**) and 24h (**D**). MCC was also calculated for NPY-emdGFP-SBP for comparison using the images shown in **Fig. 1**. *p<0.05; ****p<0.0001 by one-way ANOVA followed by posthoc Tukey t-test (for SBP-emdGFP-VSV-G 1hr Biotin: n = 21 cells from 2 independent transfections; for SBP-emdGFP-VSV-G 24h biotin: n = 28 from 3 independent transfections for phogrin-emdGFP-SBP 1hr Biotin: n = 21 cells from 2 independent transfections; for phogrin-emdGFP-SBP 24h biotin: n = 28 from 3 independent transfections, for NPY-emdGFP-SBP 1h biotin: n = 26 cells; for NPY-emdGFP-SBP 24h biotin: n = 29 cells). (**E-H**) INS-1 cells were transfected with VMAT2-emdGFP-SBP (**E**) or SgII-emdGFP-SBP (**G**) and co-stained for insulin at 1h or 24h. Insets are shown in **E’**and **G’**. The extent of colocalization with insulin was determined by Manders Correlation Coefficient (MCC) at 1h and 24h for VMAT2-emdGFP-SBP (**F**) and SgII-emdGFP-SBP (**G**). *p<0.05, ****p<0.0001 by unpaired t-test (for VMAT2-emdGFP-VSV-G 1hr biotin: n = 21 cells, 24h biotin: n=12 cells from 2 independent transfections; for SgII-emdGFP-SBP 1h biotin: n=24 cells, 24h biotin: n = 22 cells from 3 independent transfections). Data shown indicate mean ± SEM. Scale bar indicates 10 μm and 1μm for insets.

To test this hypothesis more directly, we repeated the experiments in presence of the dynamin inhibitor dynasore in INS-1 cells. Indeed, dynamin II mediates the scission of constitutive secretory vesicles at the TGN (Jones et al., 1998, Kockx et al., 2014). As a positive control, we looked at the behavior of a bona fide transmembrane marker of constitutive secretory vesicles, SBP-emdGFP-VSV-G (**Fig. S1A**) (Rivas and Moore, 1989). We first validated that SBP-emdGFP-VSV-G trafficked properly through the secretory pathway (**Fig. 3B**, **Movie S1**). Indeed, after 1h of biotin, we could observe accumulation of this marker at the plasma membrane as expected, suggesting that it is reaching the cell surface in a constitutive manner and it did not colocalize with insulin at 1h or 24h (**Fig. 3B-D**). Dynasore treatment led to accumulation of SBP-emdGFP-VSV-G in the TGN region and slowed down its budding kinetics (**Fig. 4A, B**). As a consequence, we could not detect post-TGN carriers in cells treated with dynasore post peak, whereas they were readily detectable in vehicle-treated cells (**Fig. 4A’**arrows). In contrast, dynasore treatment slowed down budding of NPY-emdGFP-SBP to a lesser extent (**Fig. 4C, D**). As a consequence, we could still observe budded SGs in presence of the inhibitor (**Fig. 4C’**). Next, we performed the same experiments with phogrin-emdGFP-SBP and VMAT2-emdGFP-SBP and observed that dynasore treatment prevented budding from the TGN of both cargoes (**Fig. 4E, F**, **Fig. S2F**). These data suggest that transmembrane SG cargoes exit the TGN within non-regulated secretory vesicles.

**Figure 4.**
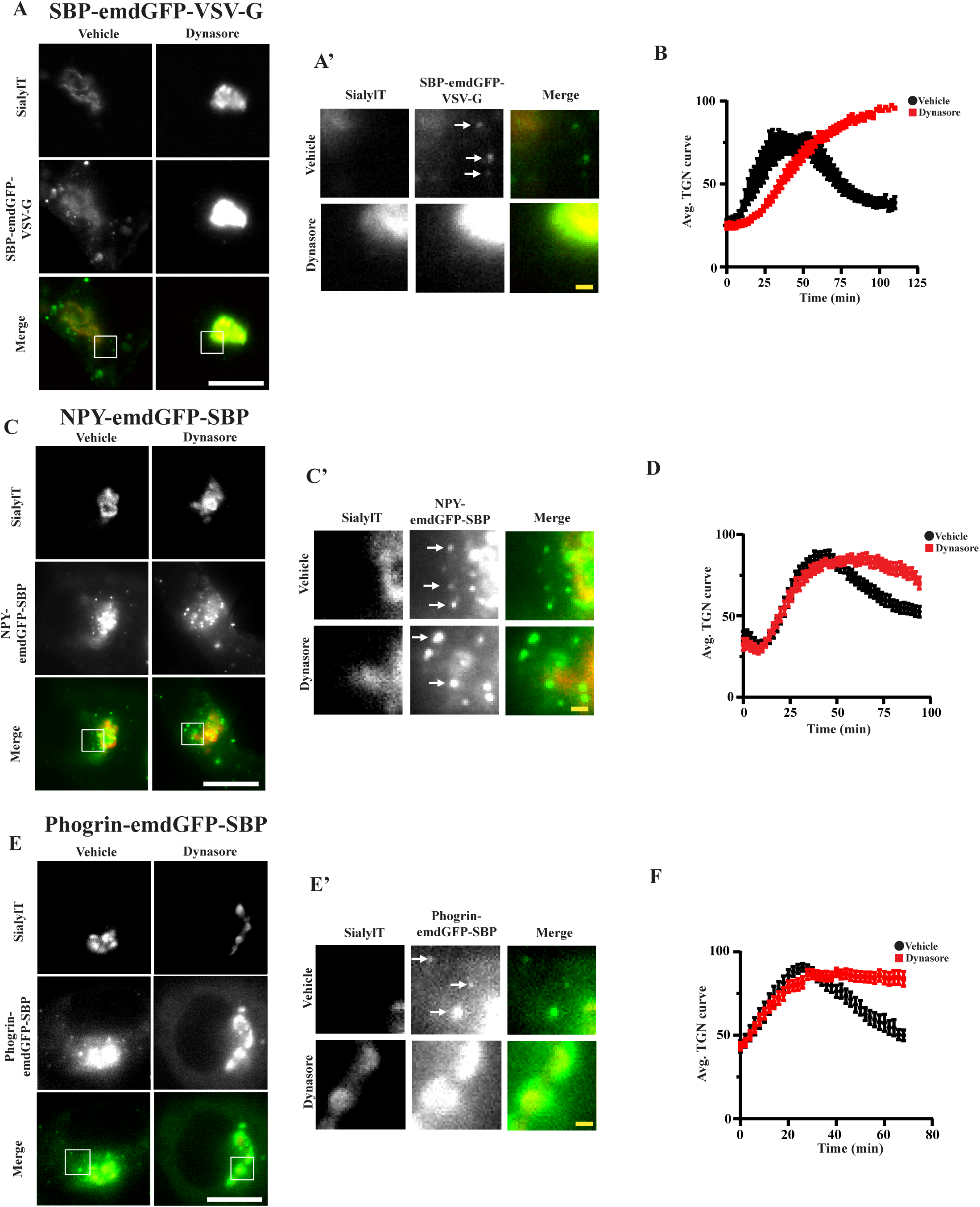
INS-1 cells transfected with the indicated cargoes were incubated with biotin in presence or absence of 80μM dynasore. Fluorescence within the TGN region was monitored as described in **Fig. 2A**. Insets are shown in **A’**, **C’** and **E’**. Arrows indicate post-TGN carriers. Average curves of TGN fluorescence are shown in (**B**), (**D**) and (**F**). (n=9 and 9 cells from 2 independent transfections for SBP-emdGFP-VSV-G vehicle and dynasore, respectively; n=19 and 20 cells from independent transfections for NPY-emdGFP-SBP vehicle and dynasore, respectively; n=7 and 12 cells from 3 independent transfections for phogrin-emdGFP-SBP vehicle and dynasore, respectively). Data shown indicate mean ± SEM. Scale bar indicates 10μm and 1μm for insets.

### Simultaneous visualization of constitutive and regulated secretory cargo budding from TGN

In order to further distinguish sorting of cargoes to the constitutive and regulated secretory pathways, we next tested whether we could follow two cargoes simultaneously. For this, we generated NPY-mCherry-SBP that we co-expressed with SgII-emdGFP-SBP, phogrin-emdGFP-SBP or SBP-emdGFP-VSV-G. To visualize budding from the TGN, we imaged our cells after 40 min of biotin treatment in order to get a wave of cargoes to the TGN. To our knowledge, this represents the first successful attempt at simultaneously imaging cargo sorting to constitutive vs regulated secretory pathways in real-time. When looking at two soluble regulated secretory cargoes, the majority of budding vesicles contained both NPY-mCherry-SBP and SgII-emdGFP-SBP (**Fig. 5A-C**, **Movie S2**). On the other hand, we observed significantly less NPY-mCherry-SBP vesicles being positive for SBP-emdGFP-VSV-G (**Fig. 5B, C**, **Movie S2**), as expected for two cargoes known to traffic to the regulated and constitutive secretory pathway, respectively. Interestingly, the proportion of NPY-mCherry-SBP vesicles also positive for phogrin-emdGFP-SBP was similar to SBP-emdGFP-VSV-G (**Fig. 5B, C**, **Movie S2**), indicating again that phogrin exits the TGN within non-regulated secretory vesicles.

**Figure 5.**
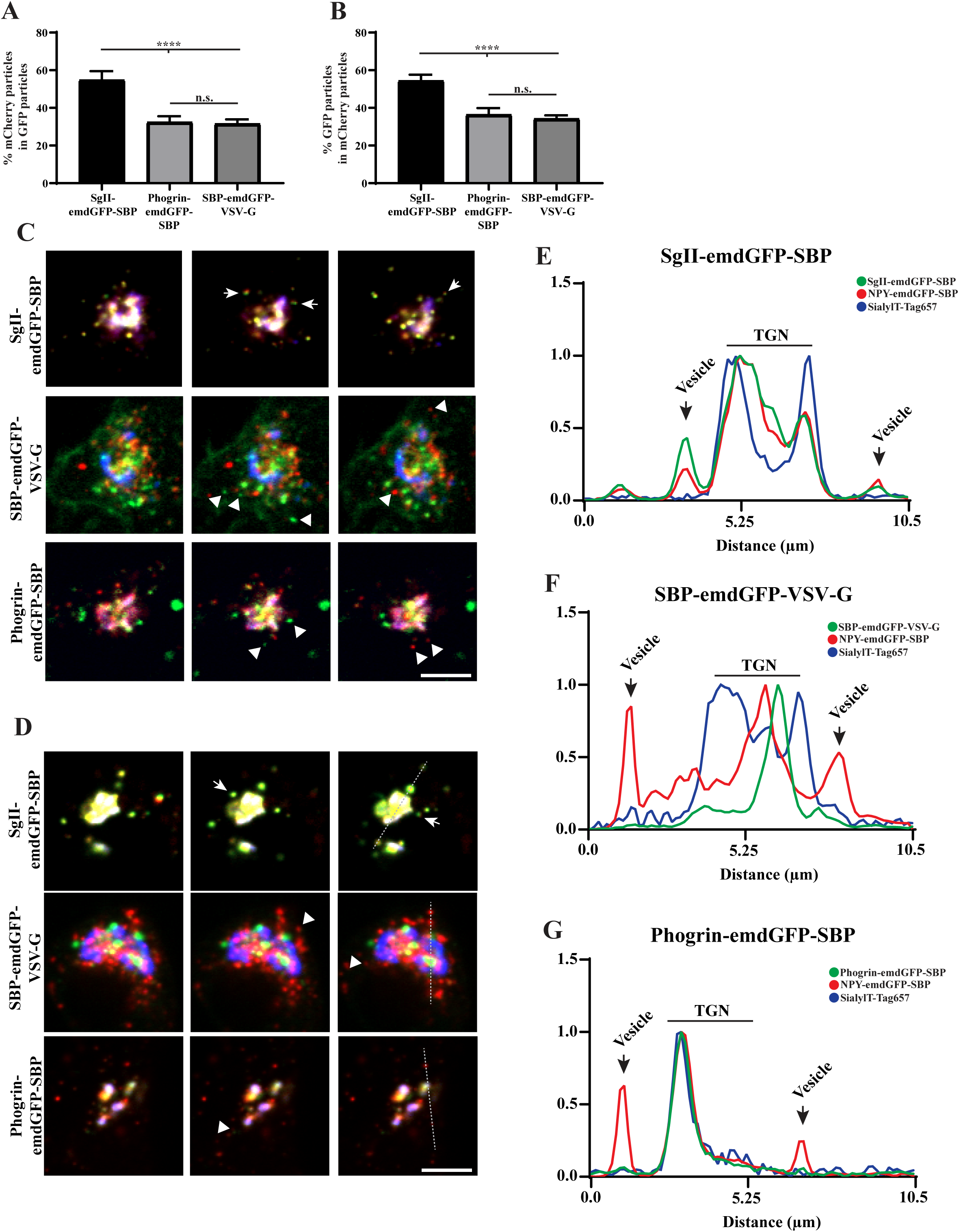
(**A-C**) INS-1 cells co-transfected with NPY-mCherry-SBP and indicated emdGFP-RUSH constructs as in **Fig. 2** and imaged every 300ms for 2min after 40min of biotin treatment. The proportion of vesicles positive for both markers was determined (**A, B**). ****p < 0.0001 (n=14 cells for SgII-emdGFP-SBP, n=12 for phogrin-emdGFP-SBP, n=19 for SBP-emdGFP-VSV-G from 2 independent transfections) by one-way ANOVA followed by posthoc Tukey t-test. Data shown indicate mean ± SEM. Consecutive frames from the movies are shown in **C**. (**D-G**) INS-1 cells co-transfected with NPY-mCherry-SBP and indicated emdGFP-RUSH constructs as in **Fig. 2** and imaged every 300ms for 2min after 40min of biotin and dynasore treatment. Consecutive frames from the movies are shown in **D**. (**E-G**) Line-scan profiles represent the fluorescence intensity across the TGN and its vicinity. Arrows indicate vesicles positive for both markers; arrowheads indicate vesicles positive for one marker. Scale bar indicates 10μm.

Finally, we repeated the same experiment in presence of dynasore to block or slow down vesicle budding from the TGN. The treatment impeded vesicular movement of all studied cargoes (**Movie S2)**, but the effect seemed more pronounced for SBP-emdGFP-VSV-G and phogrin-emdGFP-SBP, resulting in a more arrested distribution around the Golgi area. NPY-mCherry-SBP did not colocalize with these arrested vesicles, whereas it did colocalize with SgII-emdGFP-SBP (**Fig. 5D-G**). Additionally, these soluble regulated secretory cargoes seemed more dynamic. Altogether these results indicate that phogrin-emdGFP-SBP do not exit the Golgi within vesicles containing soluble regulated secretory cargoes and suggests that it gets added to SGs post-budding. Interestingly, analysis of phogrin recycling after exocytosis has revealed that it can reach an insulin-positive compartment from the plasma membrane without going through the TGN (Vo et al., 2004). Consistent with this idea, several genes typically associated with the endocytic and retrograde pathways affect SG formation (Asensio et al., 2010, Asensio et al., 2013, Sasidharan et al., 2012, Sirkis et al., 2013, Paquin et al., 2016, Topalidou et al., 2016, Zhang et al., 2017). It will be interesting to determine the exact step and molecular players controlling the recruitment of phogrin, and whether this mechanism holds true for all transmembrane SG cargoes.

## Material and Methods

### Molecular biology

The following primers were used for cloning. P1: 5’-ACCGGTCCACCATGTTAGGTAACAAGCG-3’; P2: 5’-CTTGCTCACAAGCTTCCACATTGCAGGG-3’; P3: 5’-ATGTGGAAGCTTGTGAGCAAGGGCGAGGA-3’; P4: 5’-TCTCCTGAACCTCCGAGCTCCTTGTACAGCTCGTCCATGC-3’; P5: 5’-GAGCTCGGAGGTTCAGGAGACGAGAAGACCACTGGTTG-3’; P6:5’-GAATTCTTATGGTTCACGTTGACCTTGTG-3’; P7: 5’-AGCTTTACCCATACGACGTACCAGATTACGCTGAGCT-3’; P8: 5’-CAGCGTAATCTGGTACGTCGTATGGGTAA-3’; P9: 5’-CCGTCAGATCCGCTAGCGCTACCGGTCCACCATGGGGCTACCGCTCCCG-3’; P10: 5’-CGCCCTTGCTCACTACCTTTCCTTTCGGCCTGATTCCA-3’; P11: 5’-AGGTAGTGAGCAAGGGCGAGGAGC-3’; P12: 5’-TAGGAGCTTGAGTCCTGGTTCACGTTGACCTTGTG-3’; P13: 5’-GAACCAGGACTCAAGCTCCTACCCCATCCAG-3’; P14: 5’-CCGTCAGATCCGCTAGCGCTACCGGTCCACCATGACTGAATCGAAGGCTTACC-3’; P15: 5’-AGCTCCTCGCCCTTGCTCACAAGCTTCATGTTTTCCATGGCCCGT-3’; P16: 5’-GGAGGAGGttcaGGAGAATTCaagttcaccatagtttttccac-3’; P17: 5’-GGATCCCGGGCCCGCGGTACCTTActttccaagtcggttcatc-3’; P18: 5’-ATAACCGGTAGCACTAGCGGCGGAAGCGGCGGGACAGGTGGAGTGAGCAAGGGCG AGGAG-3’; P19: 5’-ATAACCGGTGCCACCGGAGCCTCCGGTTCCGCCGCTGCCACCTGGTTCACGTTGACC TTG-3’; P20: 5’-ATATAAGCTTGTGAGCAAGGGCGAGGAG-3’; P21: 5’-ATATGAGCTCcttgtacagCTCGTCCATGCC-3’. Previous studies utilizing RUSH performed the assay using a bicistronic plasmid containing an ER hook (KDEL-streptavidin) followed by a “hookable” cargo after an IRES (Boncompain et al., 2012, Chen et al., 2017, Emperador-Melero et al., 2018). When these cargoes were tested in COS-7 cells, they behaved as expected (data not shown), but they did not in (neuro-)endocrine cells. The fusion proteins expressed at very low levels and the cells did not display any RUSH upon biotin addition. To alleviate this, we expressed the hook and cargo using two separate plasmids, which solved the issue of low expression, however, this alone did not allow for RUSH to occur due to cargo not being hooked properly. We further engineered our ER hook by introducing a linker between the streptavidin and the KDEL sequence. This modification was essential and sufficient to enable efficient hooking at the ER. All constructs used were generated using pemdGFPC3 as a backbone. STR-HA-KDEL: HA-KDEL was produced by annealing P19 and P20. The product was used in a ligation reaction with pemdGFPC3-STR cut with Sbf1 and Sac1. To generate STR-Linker-KDEL, we introduced a flexible linker by annealing P21 and P22 and ligating into STR-HA-KDEL cut with SbfI and SacI, which removed the HA. To generate NPY-emdGFP-SBP, we amplified NPY using P1 and P2, emdGFP using P3 and P4, and SBP using P5 and P6. We performed overlap extension PCR using emdGFP and SBP fragments. NPY, emdGFP-SBP, and pemdGFPC3 cut with AgeI and EcoR1 were used in a three-part Gibson reaction. To generate NPY-HA-SBP, HA was obtained by annealing P7 and P8. The annealed primers were used in a ligation reaction with NPY-emdGFP-SBP cut with HindIII and SacI, which removed emdGFP. To generate phogrin-emdGFP-SBP, phogrin’s lumenal domain up to the transmembrane domain was amplified using P9 and P9. emdGFP-SBP was amplified using P10 and P11. The remaining portion of phogrin’s lumenal domain and the transmembrane plus cytosolic domain was amplified using P12 and P13. The emdGFP-SBP and cytosolic portion of phogrin underwent overlap extension. We introduced emdGFP-SBP within the lumenal domain (after aminoacid 588). This construct was designed based on the extensive, unpublished work done in the laboratory of the late John Hutton, where a pHluorin-phogrin fusion was generated (Bauer, 2008). This construct was validated by showing that, at this particular site, the insertion of the fluorescent protein did not affect the trafficking of phogrin in INS-1 cells. To generate SgII-emdGFP-SBP, SgII was amplified using P14 and P15 and ligated pemdGFPC3-emdGFP-SBP cut with AgeI and HindIII. To generate SBP-emdGFP-VSV-G, VSVG was amplified using P16 and P17. A vector containing SBP-emdGFP was cut with EcoRI and KpnI. PCR and vector were used in a Gibson reaction to generate SBP-emdGFP-VSVG. To generate VMAT2-emdGFP-SBP, we inserted emdGFP-SBP within the first lumenal loop of VMAT2. Indeed, insertion of pHluorin at this particular site does not affect VMAT2’s trafficking (Anantharam et al., 2010). To amplify emdGFP-SBP, we used P18 and P19 and ligated into pemdGFPC3-VMAT2 cut with AgeI. To generate NPY-mCherry-SBP, we amplified mCherry using P20 and P21 and ligated within NPY-emdGFP-SBP cut with HindIII and SacI. All constructs were verified by Sanger sequencing (Quintarabio). Primers were obtained from IDT and restriction enzymes from NEB.

### Antibodies

The following primary antibodies were used in this study: mouse anti-insulin (Assady et al., 2001) (Sigma SAB4200691), dilution 1:1000; rabbit anti-SgII (Sirkis et al., 2013) (Meridian K55101R), dilution 1:2000; rat anti-HA (Hummer et al., 2017) (Sigma 12158167001), dilution 1:500. The following secondary antibodies were used: goat anti-mouse Alexa Fluor 647 (Invitrogen A32728), dilution 1:2000; goat anti-rabbit Alexa Fluor 647 (Invitrogen A21244), dilution 1:2000; goat anti-rat Alexa Fluor 647 (Invitrogen A21247), dilution 1:2000.

### Cell culture and transfection

INS-1 cells (U of Michigan) were cultured in RPMI 1640 (Genesee Scientific, Cat# 25-506H) with 10% fetal bovine serum (VWR, Cat#89510-186) and 1mM sodium pyruvate (Genesee Scientific, Cat#25-537) in 5% CO_2_ at 37C. PC12 cells (UCSF) were cultured in DMEM (Genesee Scientific, Cat#25-501) with 10% Horse Serum and 5% Cosmic Calf Serum in 5% CO_2_ at 37C. Transfections for live-imaging were performed using Fugene HD (Promega, Cat#E2311) according to the manufacturer protocol. For RUSH, a ratio of 3:1 hook to cargo ratio was determined to be ideal. Lipofectamine 2000 was used to transfect PC12 cells for secretion assays. Experiments were performed 36-48h after transfection. Cells were routinely tested for the absence of mycoplasma. For live imaging, cells were transfected in 24-well dishes (Genesee Scientific, Cat#25-107), trypsinized and replated 24h post transfection to a 35mm glass bottom imaging dish coated with PLL (Sigma-Aldrich, Cat#P2636). Cells were imaged the following day. For live imaging, INS-1 were transferred to biotin and phenol-red free RPMI (US Biological Life Sciences, Cat#R9002-01) media supplemented with 2.4mM sodium bicarbonate and 25mM HEPES. PC12 cells were imaged in phenol-red free complete media.

### Immunofluorescence

Cells were fixed in 4% paraformaldehyde in PBS for 20min, permeabilized with 0.1% Triton X-100 in PBS, incubated in PBS block buffer (2% BSA, 1% Fish Skin Gelatin, 0.02% Saponin) for 1h, incubated with primary antibodies for 1h followed by incubation with secondary antibodies for 1h.

### Secretion Assays

PC12 or INS-1 cells were seeded onto a 24-well cell culture dish coated with or without PLL, respectively. INS-1 cells were washed once in PBS, reset in low potassium KRB (138mM NaCl, 5.4mM KCl, 2.6mM MgSO_4_, 5.0mM NaHCO_3_, 10mM HEPES, 2.6mM CaCl_2_, 1.5mM glucose) for 1hr at 37C. Following an additional PBS wash, cells were incubated with low potassium KRB with 50μM biotin (Sigma-Aldrich, Cat#B4501-1G) for 3h. Cells were washed again with PBS, and treated with low or high potassium (97.8mM NaCl, 45.6mM KCl) KRB for 15min. Secreted fractions were collected. Cells were washed with ice cold PBS, and lysed in 50mM Tris, pH 7.4, 150mM NaCl, mM EDTA, 1% TX-100 with protease inhibitors (Sigma-Aldrich) and 1mM PMSF (Sigma-Aldrich). Secreted fractions and cellular lysates were prepared and fluorescence was measured using a plate reader as described previously (Hummer et al., 2017). PC12 cells were washed with PBS, incubated with complete media in presence of 50μM biotin for 1.5h at 37C. Following an additional PBS wash, cells were incubated with low or high potassium Tyrode buffer for 15min at 37C. Secreted and cellular fractions were prepared as described above.

### Image acquisition and analysis

Images for colocalization analysis were acquired using a custom-built Nikon spinning disc confocal. Images were collected with a 63× objective (Oil Plan Apo NA 1.49) and an ImageEM X2 EM-CCD camera (Hamamatsu, Japan) at a resolution of 512 × 512 pixels. Images for puncta analysis were imaged on an Evos FL Auto 2 microscope using a 63× objective (Oil Plan Apo NA 1.42, Olympus). Manders correlation coefficient was determined using the Colocalization Threshold Fiji plugin using an ROI excluding the nucleus and Golgi region. Cells were excluded from analysis for tM1 or tM2 values of 1 due to thresholding issues and cells with no positive correlation were also excluded. Live movies were imaged on an Evos FL Auto 2 in an environmentally controlled imaging chamber (37C, 5% CO2, 20% humidity). Cells were imaged every 2 min. Image processing was performed using MATLAB software. To isolate regions of interest, each frame of the Golgi-labeled channel was manually cropped around the general cell area and segmented using an intensity threshold filter. The segmented region of interest was constrained by an area filter to reduce noise, generating a binary mask of only the largest objects within the frame. The mask was then convoluted with the cargo channel to obtain the TGN signal, and the inverse of the mask was convoluted with the same channel to acquire the background signal. For figures, images were processed using ImageJ, any changes in brightness and contrast were identical between samples meant for comparison. First order decay curves were fit using GraphPad Prism version 8.0.2 for Windows, GraphPad Software, La Jolla (California). The plateau followed by a one phase decay function with a constraint of X0 begin greater than 0 was used to fit k_TGN_. The one phase decay function was used to fit k_ER_. Average curves for experiments with dynasore were made by combining the curves from all cells starting directly after biotin addition. For phogrin, due to variable RUSH starting time, we aligned the individual curves at the beginning of ER exit to generate average curves. Statistical analysis was performed using Prism. The following criteria were used to exclude cells from further analysis: 1) cell division, 2) excessive Golgi fractionation, or 3) extensive out of focus drift. Outside of these criteria, all cells were used for analysis. For two cargoes RUSH experiments, transfected cells were imaged using a spinning-disk confocal 30-40 min after biotin addition every 300 ms for 2 min. We determined the proportion of newly budded vesicles containing both cargoes using the ComDet v.0.4.2 plugin for Image J (https://github.com/ekatrukha/ComDet) using a circular ROI around the Golgi region with a diameter equals to twice the diameter of the Golgi. Proportions were obtained for each frame, and an average proportion per cell was calculated over 150 frames.

## Acknowledgments

We thank Peter Arvan for the INS-1 cell line and Dinah Loerke for help with image analysis; Str-KDEL ManII-SBP-EGFP was a gift from Franck Perez (Addgene plasmid # 65252; http://n2t.net/addgene:65252; RRID:Addgene_65252); This work was supported by American Diabetes Association grant #1-17-JDF-064 and by National Institute of General Medical Sciences grants R01 GM124035 and R15 GM116096 to CSA. We would like to thank Katharine Roth for technical support.

## References

Anantharam, A., Onoa, B., Edwards, R. H., Holz, R. W. & Axelrod, D. 2010. Localized topological changes of the plasma membrane upon exocytosis visualized by polarized TIRFM. J Cell Biol, 188, 415–28.

Arvan, P. & Halban, P. A. 2004. Sorting ourselves out: seeking consensus on trafficking in the beta-cell. Traffic, 5, 53–61.

Asensio, C. S., Sirkis, D. W. & Edwards, R. H. 2010. RNAi screen identifies a role for adaptor protein AP-3 in sorting to the regulated secretory pathway. J Cell Biol, 191, 1173–87.

Asensio, C. S., Sirkis, D. W., Maas, J. W. Jr, Egami, K., To, T. L., Brodsky, F. M., Shu, X., Cheng, Y. & Edwards, R. H. 2013. Self-assembly of VPS41 promotes sorting required for biogenesis of the regulated secretory pathway. Dev Cell, 27, 425–37.

Assady, S., Maor, G., Amit, M., Itskovitz-Eldor, J., Skorecki, K. L. & Tzukerman, M. 2001. Insulin production by human embryonic stem cells. Diabetes, 50, 1691–7.

Bauer, R. A. 2008. Characterization of sorting motifs in the dense core vesicle membrane protein phogrin. Dissertation/Thesis, ProQuest Dissertations Publishing.

Blagoveshchenskaya, A. D., Hewitt, E. W. & Cutler, D. F. 1999. A complex web of signal-dependent trafficking underlies the triorganellar distribution of P-selectin in neuroendocrine PC12 cells. J Cell Biol, 145, 1419–33.

Boncompain, G., Divoux, S., Gareil, N., De Forges, H., Lescure, A., Latreche, L., Mercanti, V., Jollivet, F., Raposo, G. & Perez, F. 2012. Synchronization of secretory protein traffic in populations of cells. Nat Methods, 9, 493–8.

Borgonovo, B., Ouwendijk, J. & Solimena, M. 2006. Biogenesis of secretory granules. Curr Opin Cell Biol, 18, 365–70.

Chen, Y., Gershlick, D. C., Park, S. Y. & Bonifacino, J. S. 2017. Segregation in the Golgi complex precedes export of endolysosomal proteins in distinct transport carriers. J Cell Biol, 216, 4141–4151.

Courel, M., Vasquez, M. S., Hook, V. Y., Mahata, S. K. & Taupenot, L. 2008. Sorting of the neuroendocrine secretory protein Secretogranin II into the regulated secretory pathway: role of N- and C-terminal alpha-helical domains. J Biol Chem, 283, 11807–22.

Dittie, A. S., Hajibagheri, N. & Tooze, S. A. 1996. The AP-1 adaptor complex binds to immature secretory granules from PC12 cells, and is regulated by ADP-ribosylation factor. J Cell Biol, 132, 523–36.

Dittie, A. S., Thomas, L., Thomas, G. & Tooze, S. A. 1997. Interaction of furin in immature secretory granules from neuroendocrine cells with the AP-1 adaptor complex is modulated by casein kinase II phosphorylation. EMBO J, 16, 4859–70.

El Meskini, R., Galano, G. J., Marx, R., Mains, R. E. & Eipper, B. A. 2001. Targeting of membrane proteins to the regulated secretory pathway in anterior pituitary endocrine cells. J Biol Chem, 276, 3384–93.

Emperador-Melero, J., Huson, V., Van Weering, J., Bollmann, C., Fischer Von Mollard, G., Toonen, R. F. & Verhage, M. 2018. Vti1a/b regulate synaptic vesicle and dense core vesicle secretion via protein sorting at the Golgi. Nat Commun, 9, 3421.

Gandasi, N. R., Vesto, K., Helou, M., Yin, P., Saras, J. & Barg, S. 2015. Survey of Red Fluorescence Proteins as Markers for Secretory Granule Exocytosis. PLoS One, 10, e0127801.

Hou, J. C., Min, L. & Pessin, J. E. 2009. Insulin granule biogenesis, trafficking and exocytosis. Vitam Horm, 80, 473–506.

Hummer, B. H., De Leeuw, N. F., Burns, C., Chen, L., Joens, M. S., Hosford, B., Fitzpatrick, J. J. & Asensio, C. S. 2017. HID-1 controls formation of large dense core vesicles by influencing cargo sorting and trans-Golgi network acidification. Mol Biol Cell, 28, 3870–3880.

Jones, S. M., Howell, K. E., Henley, J. R., Cao, H. & Mcniven, M. A. 1998. Role of dynamin in the formation of transport vesicles from the trans-Golgi network. Science, 279, 573–7.

Kockx, M., Karunakaran, D., Traini, M., Xue, J., Huang, K. Y., Nawara, D., Gaus, K., Jessup, W., Robinson, P. J. & Kritharides, L. 2014. Pharmacological inhibition of dynamin II reduces constitutive protein secretion from primary human macrophages. PLoS One, 9, e111186.

Krantz, D. E., Waites, C., Oorschot, V., Liu, Y., Wilson, R. I., Tan, P. K., Klumperman, J. & Edwards, R. H. 2000. A phosphorylation site regulates sorting of the vesicular acetylcholine transporter to dense core vesicles. J Cell Biol, 149, 379–96.

Li, H., Waites, C. L., Staal, R. G., Dobryy, Y., Park, J., Sulzer, D. L. & Edwards, R. H. 2005. Sorting of vesicular monoamine transporter 2 to the regulated secretory pathway confers the somatodendritic exocytosis of monoamines. Neuron, 48, 619–33.

Liu, Y., Schweitzer, E. S., Nirenberg, M. J., Pickel, V. M., Evans, C. J. & Edwards, R. H. 1994. Preferential localization of a vesicular monoamine transporter to dense core vesicles in PC12 cells. J Cell Biol, 127, 1419–33.

Milgram, S. L., Mains, R. E. & Eipper, B. A. 1996. Identification of routing determinants in the cytosolic domain of a secretory granule-associated integral membrane protein. J Biol Chem, 271, 17526–35.

Oprins, A., Rabouille, C., Posthuma, G., Klumperman, J., Geuze, H. J. & Slot, J. W. 2001. The ER to Golgi interface is the major concentration site of secretory proteins in the exocrine pancreatic cell. Traffic, 2, 831–8.

Paquin, N., Murata, Y., Froehlich, A., Omura, D. T., Ailion, M., Pender, C. L., Constantine-Paton, M. & Horvitz, H. R. 2016. The Conserved VPS-50 Protein Functions in Dense-Core Vesicle Maturation and Acidification and Controls Animal Behavior. Curr Biol, 26, 862–71.

Park, J. J., Koshimizu, H. & Loh, Y. P. 2009. Biogenesis and transport of secretory granules to release site in neuroendocrine cells. J Mol Neurosci, 37, 151–9.

Rivas, R. J. & Moore, H. P. 1989. Spatial segregation of the regulated and constitutive secretory pathways. J Cell Biol, 109, 51–60.

Sasidharan, N., Sumakovic, M., Hannemann, M., Hegermann, J., Liewald, J. F., Olendrowitz, C., Koenig, S., Grant, B. D., Rizzoli, S. O., Gottschalk, A. & Eimer, S. 2012. RAB-5 and RAB-10 cooperate to regulate neuropeptide release in Caenorhabditis elegans. Proc Natl Acad Sci U S A, 109, 18944–9.

Sirkis, D. W., Edwards, R. H. & Asensio, C. S. 2013. Widespread Dysregulation of Peptide Hormone Release in Mice Lacking Adaptor Protein AP-3. PLoS Genet, 9, e1003812.

Suckale, J. & Solimena, M. 2010. The insulin secretory granule as a signaling hub. Trends Endocrinol Metab, 21, 599–609.

Taraska, J. W., Perrais, D., Ohara-Imaizumi, M., Nagamatsu, S. & Almers, W. 2003. Secretory granules are recaptured largely intact after stimulated exocytosis in cultured endocrine cells. Proc Natl Acad Sci U S A, 100, 2070–5.

Tooze, S. A. 1998. Biogenesis of secretory granules in the trans-Golgi network of neuroendocrine and endocrine cells. Biochim Biophys Acta, 1404, 231–44.

Tooze, S. A. & Huttner, W. B. 1990. Cell-free protein sorting to the regulated and constitutive secretory pathways. Cell, 60, 837–47.

Topalidou, I., Cattin-Ortola, J., Pappas, A. L., Cooper, K., Merrihew, G. E., Maccoss, M. J. & Ailion, M. 2016. The EARP Complex and Its Interactor EIPR-1 Are Required for Cargo Sorting to Dense-Core Vesicles. PLoS Genet, 12, e1006074.

Torii, S., Saito, N., Kawano, A., Zhao, S., Izumi, T. & Takeuchi, T. 2005. Cytoplasmic transport signal is involved in phogrin targeting and localization to secretory granules. Traffic, 6, 1213–24.

Tsuboi, T., Zhao, C., Terakawa, S. & Rutter, G. A. 2000. Simultaneous evanescent wave imaging of insulin vesicle membrane and cargo during a single exocytotic event. Curr Biol, 10, 1307–10.

Vo, Y. P., Hutton, J. C. & Angleson, J. K. 2004. Recycling of the dense-core vesicle membrane protein phogrin in Min6 beta-cells. Biochem Biophys Res Commun, 324, 1004–10.

Weihe, E., Schafer, M. K., Erickson, J. D. & Eiden, L. E. 1994. Localization of vesicular monoamine transporter isoforms (VMAT1 and VMAT2) to endocrine cells and neurons in rat. J Mol Neurosci, 5, 149–64.

Zhang, X., Jiang, S., Mitok, K. A., Li, L., Attie, A. D. & Martin, T. F. J. 2017. BAIAP3, a C2 domain-containing Munc13 protein, controls the fate of dense-core vesicles in neuroendocrine cells. J Cell Biol, 216, 2151–2166.

