## Supplemental Figure for "Differential sorting behavior for soluble and transmembrane cargoes at the trans-Golgi network in endocrine cells"

**Supplemental Materials, Hummer et al.**

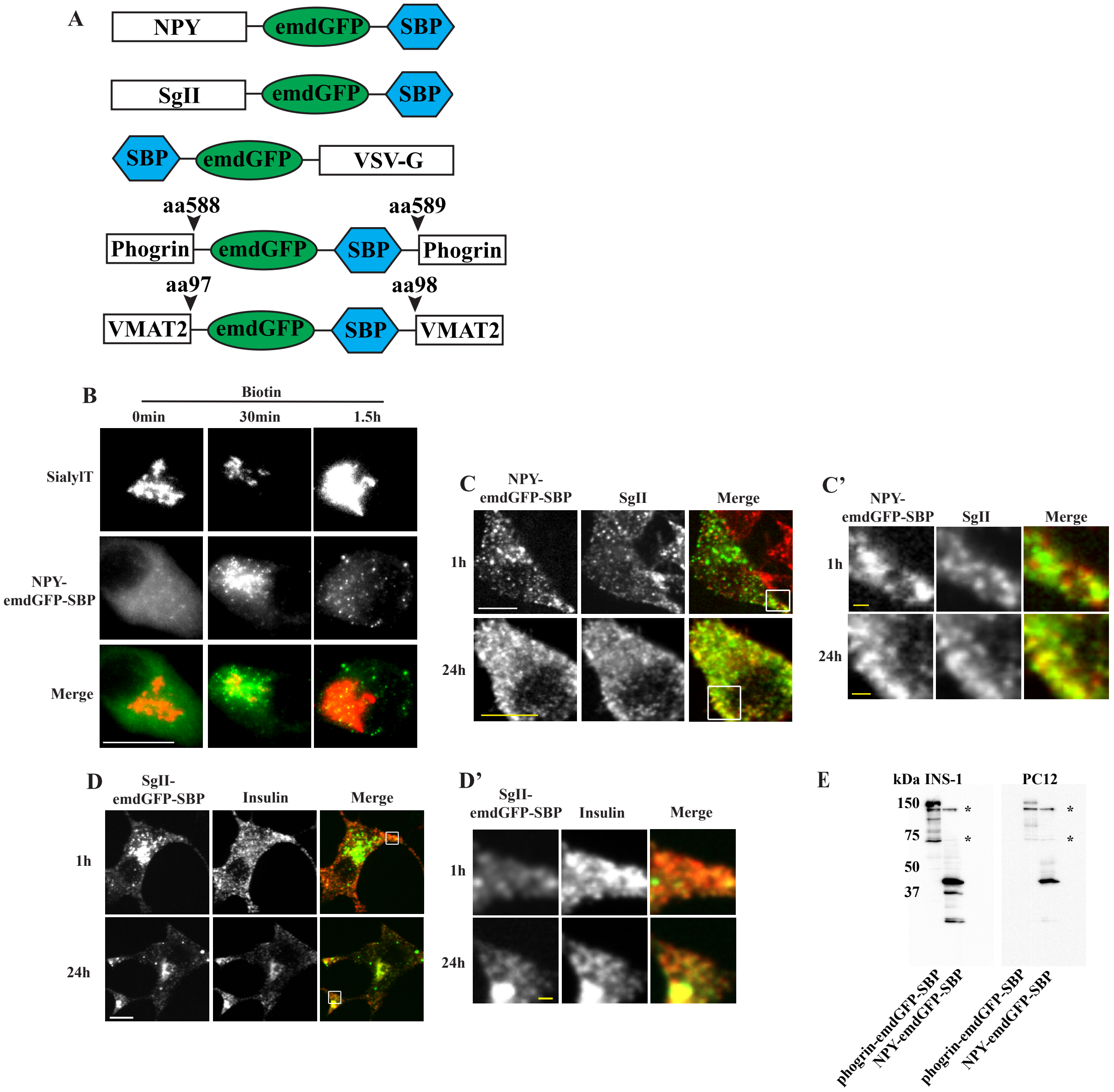


*Hummer et. al. Supplemental Figure 1*

**Figure S1**

(**A**) Diagrams depicting the insertion sites of GFP-SBP for the RUSH constructs used in this study. (**B**) PC12 cells transfected with NPY-emdGFP-SBP as in **Fig. 1**, incubated with biotin for the indicated times, fixed and imaged using a widefield epifluorescence microscope. (**C**) PC12 cells transfected with NPY-emdGFP-SBP and incubated with biotin for 1h (**B**) or 24h (**C**) and co-stained for SgII. Insets are shown in **C'**. (**D**) INS-1 cells transfected with SgII-emdGFP-SBP and incubated with biotin for 1h or 24h and co-stained with insulin. Insets are shown in **D'**. (**E**) Western blot using lysates from INS-1 or PC12 cells transfected with phogrin-emdGFP-SBP or NPY-emdGFP-SBP as indicated. Asterisks indicate non-specific bands. Scale bar indicates 10 μm and 1 μm for insets.

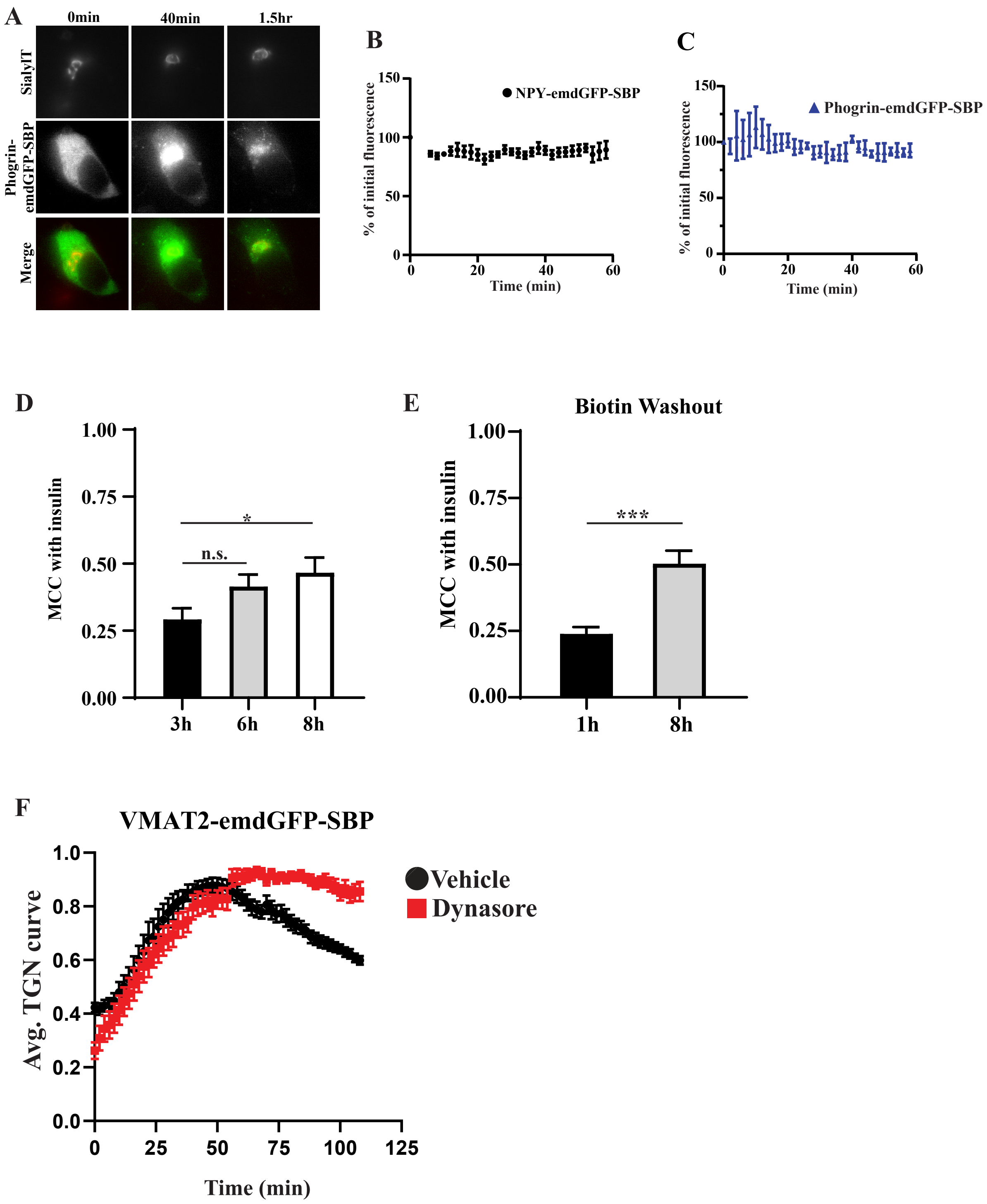


*Hummer et. al. Supplemental Figure 2*

**Figure S2**

(**A**) INS-1 cells were transfected with phogrin-emdGFP-SBP together with the TGN marker sialyltransferase-TagRFP657 and ER-hook, incubated with biotin for the indicated times, fixed and imaged using a widefield epifluorescence microscope. (**B,C**) INS-1 cells were co-transfected with indicated RUSH cargoes together with an ER-hook and imaged using an inverted widefield microscope. Cells not treated with biotin were imaged as in **Fig. 2** to measure photobleaching. n = 3 cells for NPY-emdGFP-SBP and n =2 for phogrin-emdGFP-SBP. Data shown indicate mean ± SEM. (**D**) INS-1 cells were transfected with phogrin-emdGFP-SBP and co-stained for insulin at the indicated times. The extent of colocalization with insulin was determined by Manders Correlation Coefficient (MCC) as in **Fig. 3**. ****$p<0.0001$ by one-way ANOVA followed by posthoc Tukey t-test (3hr Biotin: n = 13 cells, 6h biotin biotin: n = 12 cells, 8h biotin: n = 8 cells). (**E**) INS-1 cells were transfected as in **D**, incubated with biotin for 30min, and rinsed extensively. Cells were fixed and co-stained for insulin at 1h or 8h after the washout. The extent of colocalization was determined as in **D**. ***$p<0.0001$ (1h biotin: n = 10 cells, 8h biotin: n = 10 cells). (**F**) INS-1 cells transfected with VMAT2-emdGFP-SBP were incubated with biotin in presence or absence of 80μM dynasore. Fluorescence within the TGN region was monitored as described in **Fig. 3**. Average curves of TGN fluorescence are shown. (n=12 and 4 cells from 2 independent transfections for VMAT2-emdGFP-SBP vehicle and dynasore, respectively). Data shown indicate mean ± SEM.

**Movie S1**

INS-1 cells transfected with sialyltransferase-TagRFP657 (red), NPY-emdGFP-SBP (green), or SgII-emdGFP-SBP (green), or phogrin-emdGFP-SBP (green), or VMAT2-emdGFP-SBP (green), or SBP-emdGFP-VSV-G (green) and ER hook. Movie shown are played at 15 frames per second. Scale bar indicates 10 μm.

**Movie S2**

INS-1 cells transfected with NPY-mCherry-SBP (red), sialyltransferase-TagRFP657 (blue), SgII-emdGFP-SBP (green), or phogrin-emdGFP-SBP (green), or SBP-emdGFP-VSV-G (green), and ER hook in presence or absence of dynasore as indicated. Movie shown is 2 min and played at 10 frames per second. Movie begins 40 min after biotin addition. Scale bar indicates 5 μm.
